# BEERS2: RNA-Seq simulation through high fidelity *in silico* modeling

**DOI:** 10.1101/2023.04.21.537847

**Authors:** Thomas G. Brooks, Nicholas F. Lahens, Antonijo Mrčela, Dimitra Sarantopoulou, Soumyashant Nayak, Amruta Naik, Shaon Sengupta, Peter S. Choi, Gregory R. Grant

## Abstract

Simulation of RNA-seq reads is critical in the assessment, comparison, benchmarking, and development of bioinformatics tools. Yet the field of RNA-seq simulators has progressed little in the last decade. To address this need we have developed BEERS2, which combines a flexible and highly configurable design with detailed simulation of the entire library preparation and sequencing pipeline. BEERS2 takes input transcripts (typically fully-length mRNA transcripts with polyA tails) from either customizable input or from CAMPAREE simulated RNA samples. It produces realistic reads of these transcripts as FASTQ, SAM, or BAM formats with the SAM or BAM formats containing the true alignment to the reference genome. It also produces true transcript-level quantification values. BEERS2 combines a flexible and highly configurable design with detailed simulation of the entire library preparation and sequencing pipeline and is designed to include the effects of polyA selection and RiboZero for ribosomal depletion, hexamer priming sequence biases, GC-content biases in PCR amplification, barcode read errors, and errors during PCR amplification. These characteristics combine to make BEERS2 the most complete simulation of RNA-seq to date. Finally, we demonstrate the use of BEERS2 by measuring the effect of several settings on the popular Salmon pseudoalignment algorithm.

## Introduction

Since the introduction of RNA-Seq circa 2010, there has been a tremendous proliferation of bioinformatics tools published for every stage of data processing and analysis, for example over a dozen differential expression methods exist^1^. Such a landscape of tools naturally calls for unbiased benchmarking studies to compare and evaluate their performances and to develop best practice guidelines. Benchmarking studies are most informative when there is data for which the ground truth is known. In the case of real world omics data, the ground truth is impossible to determine without the use of the same tools being evaluated, and real-world data is inherently limited for evaluating metrics^2^.

Simulation is a popular solution, however, the further one goes downstream in the analysis pipeline of RNA-Seq, the more realistic the simulation must be. Benchmarking genome alignment with simulated data only requires simulating realistic reads because genome aligners only work on one read at a time. However, benchmarking the downstream tasks of quantification and normalization require simulating entire realistic samples of reads because these steps combine the information across reads, and modern methods like pseudoalignment^3, 4^ can combine traditional alignment and quantification into a single step. Therefore, there is a strong need for sophisticated RNA-Seq simulation, and current simulators are too simplistic^5^.

Developing read-level simulated data from scratch is labor-intensive and depending on the specific application may not be necessary. Indeed, much of the benchmarking of RNA-seq tasks in the literature^6-9^ involves directly simulating the final normalized spreadsheet of gene, or transcript, level quantifications without ever simulating reads. However, this spreadsheet level simulation cannot fully capture the difficulties of handling real reads^10^ and read level simulation is necessary to cover the full spectrum of RNA-Seq analysis methods. Both alignment and quantification, for example, require read-level data.

When embarking on a benchmarking study with simulated read-level data, it is necessary to budget a fair amount of time, effort and thought into generating the simulated data. Otherwise, the results are only as meaningful as the data is realistic and it is very easy to oversimplify when simulating biology. In this work we have provided powerful tools to achieve reads and samples of reads that are as realistic as possible. But these are not push-button applications, they are aids to science that must be employed with expert knowledge.

We built the original RNA-Seq read-level simulator in 2011 called BEERS^11^ specifically to benchmark alignment. However, in the years that followed, we realized that BEERS was being used to benchmark all stages of the processing and analysis pipeline, which was far beyond its intended purpose^12-14^. To this end, we present BEERS2, in which we have endeavored to model every step of the library prep pipeline *in silico*.

Simulating RNA-Seq data for general purpose involves first simulating “expression” and then simulating “sequencing”. Given the complexity of both of these steps, we have separated them into two different applications, reflecting the separation of ‘expression’ and ‘measurements’ models of RNA-seq^15^. The expression simulator is called CAMPAREE^16^ and it generates full length RNA molecules from a diploid genome inferred separately for each sample. CAMPAREE takes real RNA-Seq data as input, from which it infers variants and expression levels and generates diploid data as the ground truth.

BEERS2 takes the set of “molecules” output by CAMPAREE as input, but it can just as easily take as input any properly formatted set of molecules, which are, typically, full length RNA, but can be RNA molecules of any sort, see Figure 1 a. BEERS2 transforms these input molecules into RNA-Seq reads, by mimicking one of the various library prep protocols followed by sequencing. Briefly, this involves ribosomal depletion, fragmentation, various PCR steps, labeling, and then sequencing by synthesis, which involves flowcell hybridization and bridge amplification, and base assessment - see Figure 1 b. Errors and other features of sequencing are incurred at the various steps that they occur during library prep. The system is modular, so that modeling new library prep protocols or implementing changes to library preps requires only local changes to the modules for the steps that have been affected. For example, polyA selection can be easily swapped out for a different ribosomal depletion protocol without changing other steps in library prep simulation. Further, this modular nature also allows for isolating specific steps to investigate the effects of their specific biases on the analysis under question.

**Figure 1.**
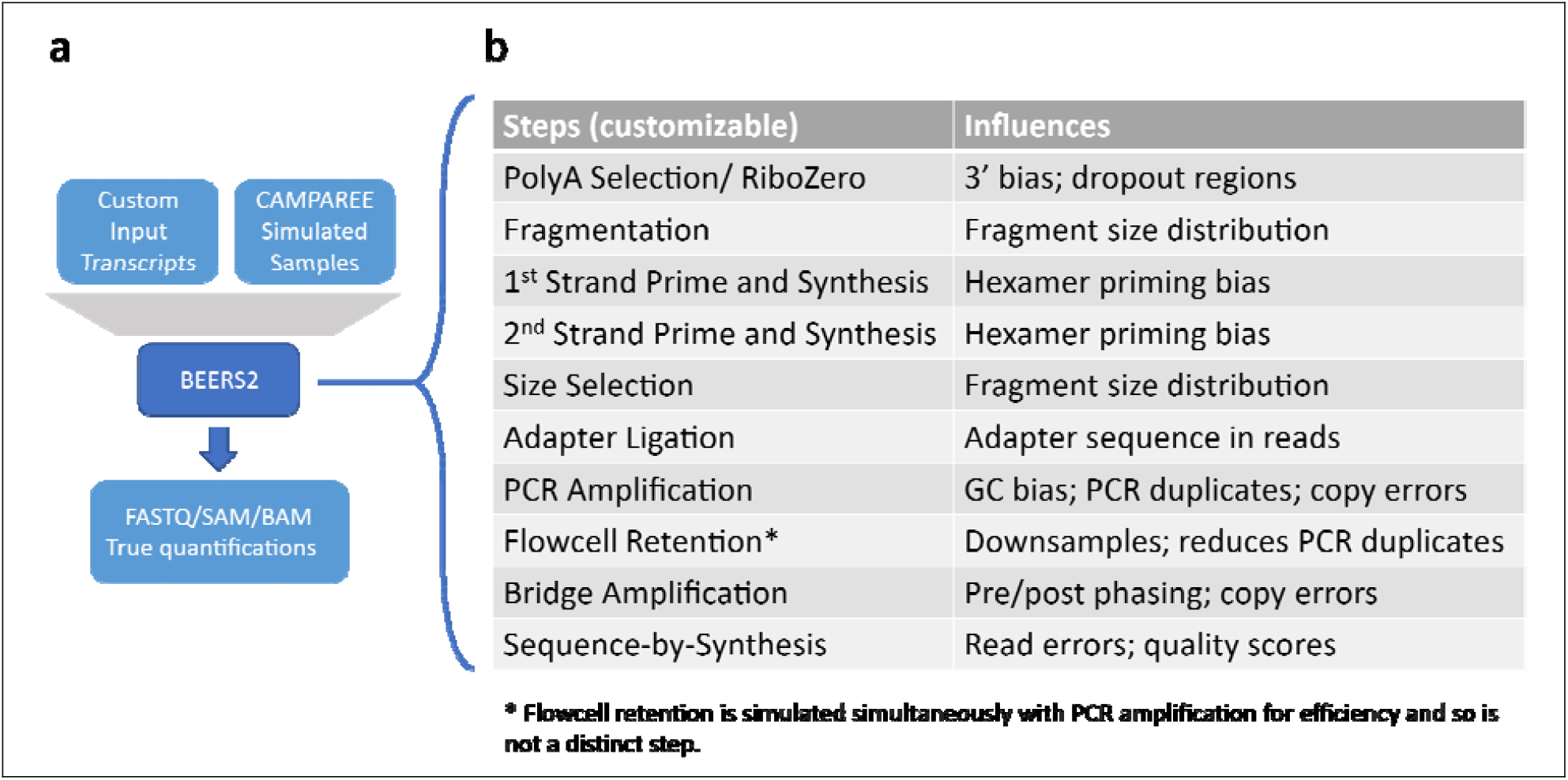
BEERS2 Overview. (a) BEERS2 is a customizable pipeline which processes input transcripts (typically fully-length mRNA transcripts with polyA tails) from either customizable input or from CAMPAREE simulated RNA samples. It produces realistic reads of these transcripts as FASTQ, SAM, or BAM formats with the SAM or BAM formats containing the true alignment to the reference genome. It also produces true transcript-level quantification values. (b) The default pipeline simulates the Illumina Stranded mRNA library preparation process, followed by sequencing. Each step influences one or more aspects of the output sample through configurable parameters.

## Methods

BEERS2 is developed in the Python v3.11 language and uses the Snakemake^17^ pipeline library v7.24.0 for parallelization and reproducibility. Since BEERS2 simulates every individual molecule through the entire library preparation and sequencing process, realistic BEERS2 executions require significant compute resources. Our Snakemake pipeline can be configured to work on many common cluster compute environments, but to further ease the process of installing the necessary software and configuring it to work on a particular cluster, we prepared a stack template which can be easily deployed in any AWS account. The deployed stack will contain both the software and hardware necessary to execute the BEERS2 pipeline, and researchers can start the execution immediately after providing the required input for BEERS2 (Figure S 1).

The modular design allows mixing and matching library pipeline steps through the configuration file. For example, a DNA-seq pipeline could be simulated by reusing the shared aspects of the RNA-seq pipeline (such as fragmentation, size selection, and PCR amplification) and dropping the distinct steps (such as polyA-tail selection). This allows BEERS2 to keep up with ever-changing protocols.

BEERS2 includes modules simulating the individual steps of the two standard RNA library preps, Illumina Stranded mRNA and Illumina Stranded Total RNA, followed by simulated sequencing by an Illumina sequencing machine. The individual steps are modeled as closely as possible to the real process. First, input molecules are selected for having a polyA tail, where the selection criteria consist of configurable acceptance probabilities as a function of tail length. This step also introduces an optional 3’ bias by modelling truncated molecules by breaking bonds between bases with uniform probability. Alternatively, the RiboZero selection module computes sequence-similarity to a population of rRNA reference oligos and degrades molecules at sites with probability dependent upon the sequence-similarity.

Next, the fragmentation step breaks molecules by default uniformly, meaning that every bond between base pairs is equally likely to break, or with a configurable positional and fragment length bias, see Supplemental Methods. Afterwards, the hexamer priming and cDNA synthesis is simulated. Hexamers are modelled as first selecting a random number of primers to bind (according to a binomial distribution with n equal to the number of potential binding sites), and then all possible binding locations are weighted according to a configurable hexamer bias model to simulate the known hexamer sequence nonuniformity^18^. Finally, the specified number of binding hexamers are then placed on random locations according to the weights and the 5’ most hexamer is used to synthesize the cDNA. Therefore, at synthesis of both cDNA strands, some fraction of end sequence may be lost if no hexamer binds at the very end of the fragment.

After cDNA is generated, the molecules are size selected with a configurable window retention probability as a function of molecule length. Then, adapters are ligated to either end. Finally, PCR amplification is modelled. Since most resulting molecules do not end up being sequenced, resulting from the process of diluting and loading samples onto an Illumina flow cell, we model the subsequent down sampling at the same time as PCR amplification. This enables efficient modelling with many rounds of PCR amplification without computing molecules that do not end up being retained. Appropriate retention rates for this step can be chosen from the known PCR cycle count with the desired PCR dupe rate, see Supplemental Methods.

Moreover, PCR amplification can have configurable copy errors (insertions, deletions, and single base errors) during each round. Errors in early rounds may therefore propagate to multiple sequenced PCR duplicates. PCR amplification also allows a configurable GC bias, with PCR amplification failing with a rate depending upon the overall GC content of the molecule.

After PCR amplification with down-sampling, molecules are run through a simulated sequencer. Molecules are bridge amplified, including single base copy errors in the amplification. This process is approximated by storing the total counts of each of A/C/G/T for each position in the molecule. Then, sequencing by synthesis is simulated by estimating fluorescence from each base pair, including phasing, followed by base-calling and quality scores derived from the process used by Illumina’s Bustard algorithm as described in^19^.

## Results

BEERS2 generates compellingly realistic RNA-seq reads under varying sequencing conditions, see Figure 2. We provide two example applications of BEERS2 to demonstrate its use in providing actionable recommendations in RNA-seq analysis through benchmarking. First, we focus on evaluating the performance of a single pseudoaligner called Salmon^4^ in order to provide quick and simple applications. To properly perform a full-scale benchmarking comparison of all tools requires a full-length article. Salmon quantifies expression at the transcript level using an RNA-Seq pseudo-alignment approach. Our previous benchmarking study found that Salmon was a top transcript-level quantifier^20^, which is the motivation to evaluate its performance further.

**Figure 2.**
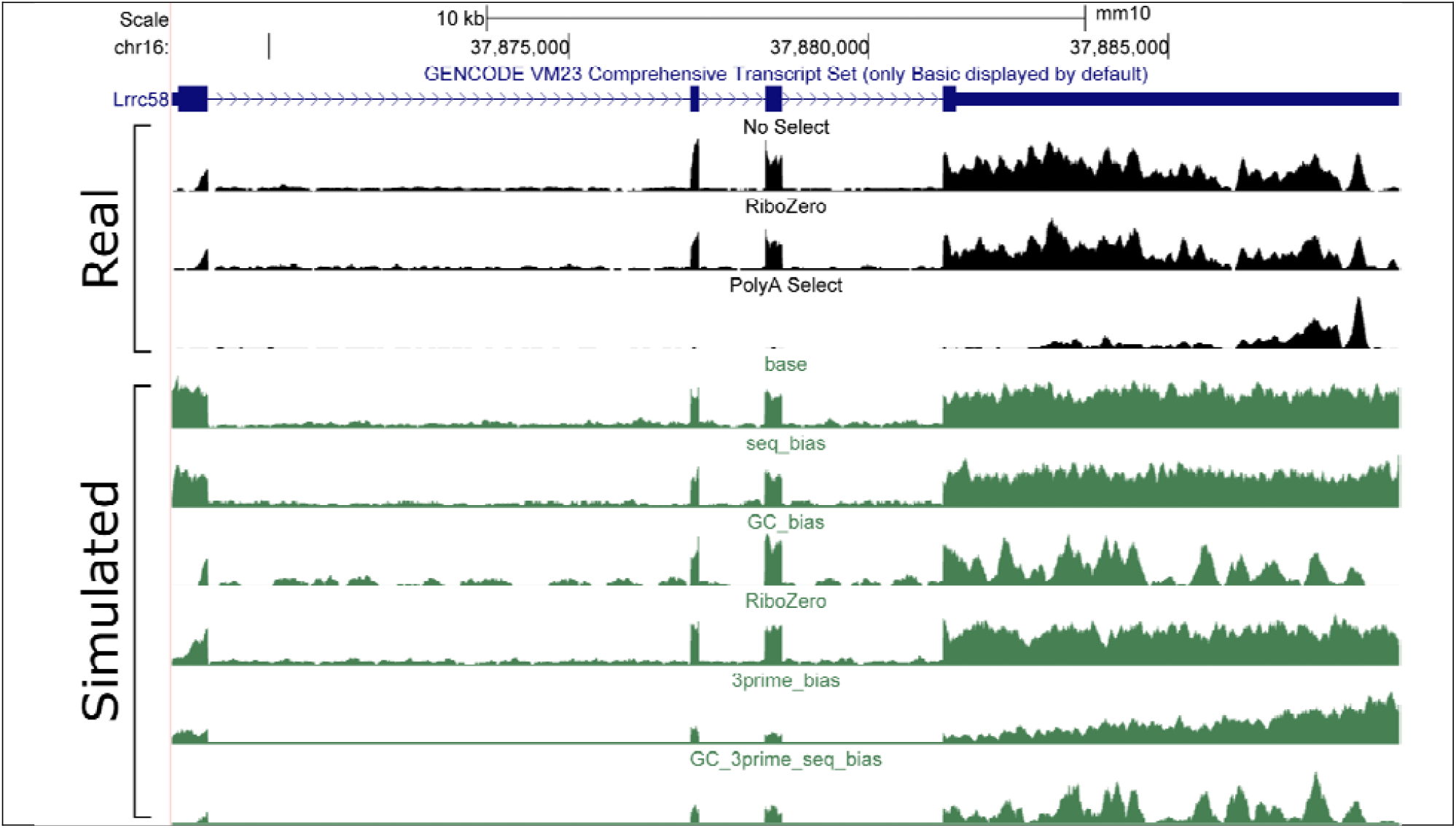
Coverage Tracks of Real and Simulated Data. UCSC Genome Browser tracks of the coverage on the gene Lrrc58 from (in black) three real mouse lung RNA-seq datasets with polyA selection, RiboZero selection, and no selection step. Below, in green, six simulated datasets under varying configurations: ideal data (base), hexamer priming sequence bias (seq_bias), GC bias in PCR amplification (GC_bias), RiboZero sequence-specific bias (RiboZero), 3’ bias from polyA selection (3prime_bias), and all three biases of a typical polyA run (GC_3prime_seq_bias). The last dataset represents the most realistic scenario. Lrrc58 gene model (blue) included at the top of the figure.

### Application 1: Benchmark of bias correction in Salmon

Salmon provides options to correct for three known bias factors in RNA-seq data: position of the fragment relative to the start and end of the transcript (Pos); base sequence near the start and end of the fragment (Seq); and GC content of the fragment (GC). Biases related to these factors can be simulated in BEERS2 by including Poly-A selection with 3’ bias (for Pos bias), primer sequence bias (for Seq bias), and PCR amplification GC bias (for GC bias). This provides an ideal test-case for BEERS2’s bias support by quantifying how Salmon’s correction options affect its accuracy in quantifying transcripts and genes under varying levels of bias. Pseudoalignment based methods such as Salmon require read level RNA-Seq benchmarking data such as provided by BEERS2.

CAMPAREE^16^ was used to simulate eight mouse liver samples based on eight real samples from a previous study^21^. Simulated sequencing of these samples was performed with BEERS2 under seven conditions with varying levels of bias:

- With no bias factors,
- With medium or high 3’ bias from polyA selection,
- With medium or high GC bias from PCR amplification,
- With primer sequence bias, or
- With high levels of all three biases (All Bias).

Salmon was then run on the FASTQ files generated by BEERS2 for each of the eight samples under each of the eight combinations of the possible bias correction settings (with/without Pos correction, with/without Seq correction, with/without GC correction), see Figure S 2. Salmon quantified TPM values were then compared to the true TPM values and Salmon read counts were compared to the true read counts. Concordance was assessed by the mean absolute relative difference (MARD) of these values, which is defined as the mean (across all transcripts) of ARD = |true – inferred| / |true + inferred|, or in other words the difference of quantified and true values divided by their sum. Lower values of MARD indicate higher mean concordance of quantified and true values. The denominator also acts as a normalization factor putting low and high expressed genes on a comparable scale.

First, we consider the TPM values, see Figure 3 a. When no biases are simulated, read depth is still not uniform primarily due to edge effects of fragmentation and size selection, see Figure S 2 a. Not surprisingly, applying the Pos correction improves accuracy only slightly. Applying either GC or Seq correction shows no improvement, and in fact slightly impaired accuracy unless Pos correction is also applied simultaneously.

**Figure 3.**
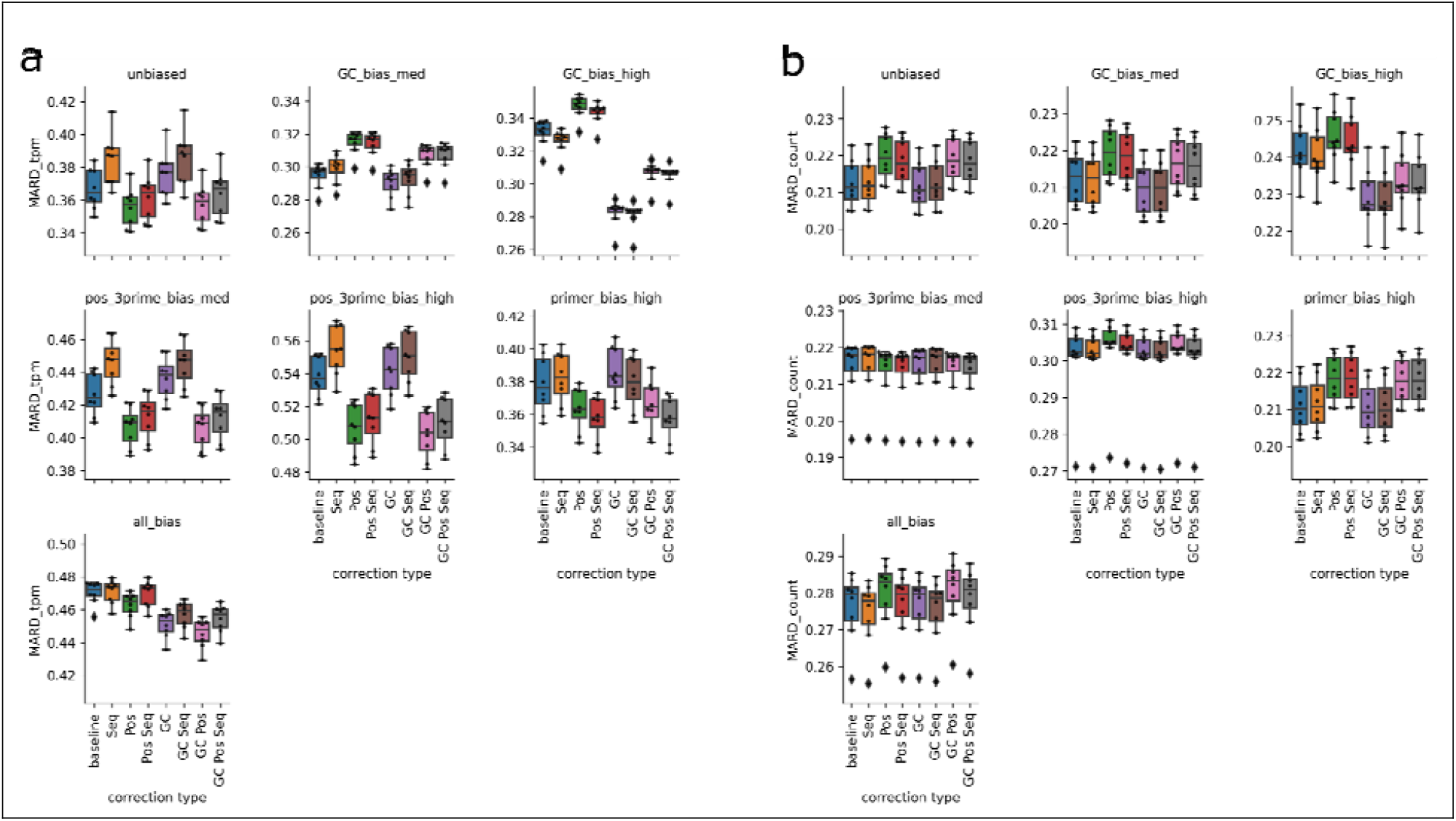
Salmon Accuracy with Bias Correction. We generated BEERS2 samples with varying levels of 3’ bias from PolyA-selection, GC bias from PCR amplification, and start and end sequence bias from primer bias for each of eight samples. Then we ran Salmon with each combination of bias correction options for fragment start/end sequence (Seq), position within transcript (Pos), and fragment GC content (GC). MARD computes the mean value of the difference of true and estimated values over their sum. Lower MARD indicate more accurate estimates. MARD values vary between samples considerably due to the relative difficulty of quantification under different bias conditions and read counts. (a) MARD of TPM values. (b) MARD of read count values. Boxes show quartiles.

As expected, when GC bias alone was present, GC correction helped performance and with 3’ bias alone, positional correction resulted in substantial improvement. Meanwhile with primer bias alone, the Seq correction improved performance but primarily only when Pos correction was also enabled.

When all three biases were included, GC and Pos correction showed substantial improvements in quantification while Seq impaired it slightly.

Next, we considered quantified read count values instead of TPM values, see Figure 3 b. Effects were largely similar but smaller than in TPM quantification with almost no changes in read count quantification on the run with all biases enabled. Read counts are overall more accurate than TPM and so may not have as much room to improve. Surprisingly, Pos correction slightly impaired instead of improved read count values when no bias or only primer bias were included, in opposite direction to the TPM values.

Comparing TPM and count values to truth by Spearman correlation instead of MARD yielded a similar picture, see Figure S 3, except that Pos correction impaired, rather than improved, quantifications when no 3’ bias was present.

Since counts were mostly unaffected by bias correction, we conclude that Salmon’s correction factors are largely affecting TPM by influencing the estimated effective length of the transcript rather than by influencing where reads counts are assigned in the expectation maximization step. This may indicate that bias correction gives limited improvement in determining which isoform an ambiguous read originates from. This is unexpected as, for example, 3’ bias effects could help disambiguate reads between isoforms with different 3’ transcription stop sites even if the read occurs far from the 3’ end.

### Application 2: Inclusion of pre-mRNA in Salmon index

Salmon indices are generated using a FASTA file containing the target transcripts and their sequences which should be quantified. These can be prepared by the user as they wish, but one common source is to use Ensembl’s cDNA files, which do not include pre-mRNA. The official Salmon *Getting Started* guide uses this file for Arabidopsis thaliana as an example. Since real RNA-seq samples invariably include intronic expression, we investigated whether Salmon’s performance would improve when including pre-mRNA in the index files. One common application of pre-mRNA quantification is RNA velocity and the preparation of indexes for that purpose in single-cell RNA-seq has been previously benchmarked^22^ while we focus instead on the impacts on mature mRNA quantification in bulk RNA-seq.

CAMPAREE generates pre-mRNA signal at realistic levels, so we used the same BEERS2-generated data from Application 1 above to assess the accuracy of Salmon with and without pre-mRNA in the index. For each gene, we added one pre-mRNA isoform to the Salmon index which contained the full genomic stretch from first to last base of all isoforms of the gene.

We compared accuracy by comparing true and estimated CPM values. Inclusion of pre-mRNA in the index slightly improved Salmon’s performance in quantifying mature mRNA among genes whose expression was at least 10% pre-mRNA (MARD 0.24 without pre-mRNA in the index versus 0.22 with; Spearman correlation of 0.935 without pre-mRNA and 0.938 with). In the genes with less than 10% pre-mRNA, including pre-mRNA in the index resulted in slightly worse performance (MARD of 0.256 without and 0.276 with pre-mRNA; Spearman correlation of 0.904 without and 0.892 with pre-mRNA).

Lastly, running Salmon with bootstrapped outputs to estimate technical variance, we noted that the relative quantification range (difference of max to min bootstrap values, divided by the mean bootstrap value) correlated modestly with ARD (Spearman correlation of 0.31 in all highly expressed transcripts and 0.39 in genes with at least 10% pre-mRNA expression). However, genes whose quantification regressed after inclusion of pre-mRNA often showed low technical variance as estimated by the Salmon bootstrap despite having large errors compared to the true simulated counts. This indicates that the difficulties in quantification are poorly captured by the bootstrap process.

We then reran all these comparisons on BEERS2 data with all biases enabled. Results were similar except that in all comparisons, inclusion of pre-mRNA made slight regressions in quantification accuracy, even in mRNA transcripts of genes with at least 10% pre-mRNA (MARD of 0.668 without and 0.700 with, and Spearman correlation of 0.799 without and 0.781 with).

We inspected genes whose quantification was most sensitive to the inclusion of pre-mRNA in the index and identified that these genes primarily displayed one of the following challenging characteristics: a large number of pseudo-genes (e.g., Rpl36a and other ribosomal proteins); annotated retained intron events (e.g., Slc35b2); overlap with an exon or intron of another gene on the same strand (e.g., Eno1b); or large overlap between an existing transcript and the included pre-mRNA (e.g., Mrpl41). However, these characteristics were common both in genes that saw improvements and those that saw regressions in quantification, indicating that they are markers of variability and not systematically affected by inclusion of the pre-mRNA index.

We conclude that inclusion of pre-mRNA in Salmon indexes has a relatively small effect and may hurt more mature mRNA quantifications than it improves. This leads to the recommendation to avoid pre-mRNA inclusion unless directly interested in pre-mRNA quantifications.

## Discussion

We demonstrate that the detailed simulation of molecule-level sequencing through BEERS2 provides valuable insights to RNA-seq analysis. In particular, it allows us to separate out individual aspects of the library preparation and sequencing to assess the impact of these steps on quantification.

In combination, CAMPAREE and BEERS2 allow simulations with substantially more detailed and configurable values. CAMPAREE includes sample-dependent features such as indel and single-nucleotide variants that are then consistent across all reads from a location, as well as a realistic distribution of reads including allele-specific expression. However, BEERS2 also remains flexible enough to accept custom input from sources other than CAMPAREE or to combine CAMPAREE-generated data and custom spike-in data. This makes BEERS2 highly adaptable for future benchmarking designs. BEERS2 allows configuration of every step of a realistic library preparation, allowing the effect of specific but known variables to be quantified.

In addition to earlier versions of BEERS, there are some existing RNA-seq read-level simulators. Polyester^23^ supports configurable fragment GC bias and differential expression; BEERS2 also supports differential expression if the user prepares appropriate input transcript samples with differential expression already present. The ASimulatoR^24^ project adds alternative-splice configuration to Polyester. The RSEM simulator^25^ allows for modelling position bias such as from polyA selection. The Flux Simulator^26^ can simulate GC bias in PCR amplification, hexamer priming sequence bias, and a 3’ bias due to poly-dT primers but not due to polyA selection. SimBA^27^ adds genomic mutations onto the Flux Simulator. CuReSim^28^ and RandomReads^29^ generate reads with configurable errors but no simulated biases. PBSIM^30^ can simulate long reads from Pacific Biosciences instruments. The rlsim^31^ package allows simulating with hexamer priming bias, and PCR amplification with GC and length biases. Thus, BEERS2 is the first simulator which models all four of the PolyA selection, RiboZero, GC, and primer biases. In addition, it provides several more advanced features: errors in PCR that propagate realistically to multiple descendant molecules; and read errors in molecule barcodes leading to demultiplexing difficulties. When used with CAMPAREE, BEERS2 also supports sample-dependent genomic variants and allele-specific diploid expression. All these factors contribute to creating more realistic simulated samples while retaining knowledge of the ground truth. Furthermore, BEERS2’s modular design makes it flexible to keep up with the fast-evolving technology of high throughput sequencing.

One limitation of the BEERS2 model is that values are configured according to how each individual step behaves. However, real data is always the result of many steps done together. Therefore, configuration values do not directly correspond to measured values and so users may have difficulty determining parameters that give realistic values for their use-case. While we provide example values as a starting point, we recommend users to experiment on small datasets to find values that fit their needs before running full simulations. BEERS2 is available for use from http://github.com/itmat/BEERS2 under a GPLv3 license. The code used for example benchmarks is available at https://github.com/tgbrooks/BEERS2_salmon_benchmark.

## Supporting information

Supplemental Manuscript

## Acknowledgements

We would like to thank Cris Lawrence for her programming expertise, Kaitlyn Forrest for processing samples for sequencing, and Jonathan Schug for insightful discussions of about sequencing.

## Data Availability

Input data used for the example applications have been previously published on the Gene Expression Omnibus repository under accession GSE98562. The generated simulated data are available for use at http://bioinf.itmat.upenn.edu/BEERS2/paper/index.html.

## Code Availability

The code for the BEERS2 simulator is available under an open-source GPLv3 license at https://github.com/itmat/BEERS2. Code used for generating and analyzing the sample application of BEERS2 is available at https://github.com/tgbrooks/BEERS2_salmon_benchmark.

