## Supplemental Manuscript for "BEERS2: RNA-Seq simulation through high fidelity *in silico* modeling"

### Supplemental Methods

#### Pseudoalignment and quantification

Salmon v1.9.0 was run on the generated FASTQ files with the following command:

salmon quant -i $INDEX -g $GTF_FILE -l A --softclip --softclipOverhangs -p 6

-1 R1.fastq -2 R2.fastq -o $OUTPUT_FOLDER –numBootstraps 50

Additionally, all combinations of the bias correction options --gcBias, --posBias, and --seqBias were run. The Salmon index was generated by:

salmon index --threads 12 -i index -t Mus_musculus.GRCm38.Ensembl.v93.cdna_ncrna_dna.fa.gz --decoy decoys.txt

Here, decoys was set to the GRCm38 DNA chromosomes. The indexed FASTA file was generated by concatenating the Ensembl GRCm38 v93 cDNA and ncRNA files along with the decoys.

#### Fragmentation step

The default fragmentation method is ‘uniform’ which fragments molecules between each base with equal probability. However, this creates a distribution that heavily favors very short and very long molecules (specifically, for all parameter settings, the most common fragment length is either 1 base or the full transcript length). After size selection, these still give reasonable fragment distributions, but we further include a configurable alternative fragmentation algorithm. Under the ‘beta’ method, fragmentation be biased towards certain positions in a fragment (e.g., near the center) by giving the parameters of a Beta distribution, which gives a probability distribution of the fragmentation position as a value 0-1 of the scaled position in the molecule. Furthermore, the rate of fragmentation can be controlled as a function of the length of the molecule. The ‘uniform’ method is equivalent to the ‘beta’ method with beta parameters A = 1, B = 1, and an exponent of 1(though ‘uniform’ fragmentation is faster to simulate than this). The ‘beta’ method then proceeds by starting with the full input molecule, determining the time until a break happens as an exponential distribution. If that time is less than the simulated fragmentation time, the molecule is then fragmentated at a location chosen by the Beta distribution. The process is then repeated on each of the resulting fragments with the fragmentation time set to the remaining time after the first break and with the new fragment’s length used to compute the rate of fragmentation.

The exact fragmentation behavior in true library prep is difficult to estimate since the end results of sequencing include many steps all at once that influence the fragments. In particular, hexamer priming, and size selection steps will alter the distribution of observed fragment end sites and lengths. Experiments design specifically to address this unknown factor would be of significant benefit for the accurate simulation of RNA-seq data.

#### PCR amplification

The PCR amplification step naïvely generates many molecules, typically about 2^14^ as many as it started with. However, most of these molecules will never be sequenced, so the PCR amplification step includes a down sampling option that reduces computational work considerably by only generating the molecules that survive down sampling. The PCR amplification step first selects how many of the descendent molecules would survive the down sampling and then, at each step of PCR amplification, assigns those surviving descendants to either the PCR copy or the original. If any molecule (copy or original) has no surviving descendants, then it is dropped from processing, thereby saving considerable compute effort as none of its descendants need to be created only to be immediately discarded.

The parameter controlling this is called retention_percentage. It can be chosen to match a real sample by examining the PCR dupe rate, for example, suppose 35% of deduped reads had one or more dupes. With a target dupe rate, in Python using the scipy library, this parameter should be chosen so that for N PCR cycles:

import scipy.stats

at_least_two = scipy.stats.binom(p=retention_percentage / 100, n=2**N).sf(1)

at_least_one = scipy.stats.binom(p=retention_percentage / 100, n=2**N).sf(0)

dupe_rate = at_least_two / at_least_one

by choosing retention_percentage to get the desired dupe_rate. For example, with N = 10 PCR cycles, retention_percentage of 0.08% gives a 35.4% dupe rate. Note that the GC bias can also induce lower PCR dupe rates by separately causing PCR amplification to fail, so if GC bias is included, this value will need to be increased.

PCR amplification steps can also accrue configurable amounts of copy errors, insertions, or deletions. If this is the case, the GC bias simulation will use the GC content of the original molecule for all daughter molecules, for the sake of efficiency, since the total GC effect of PCR copy errors, insertions, and deletions is likely to be negligible.

### Supplemental Figures

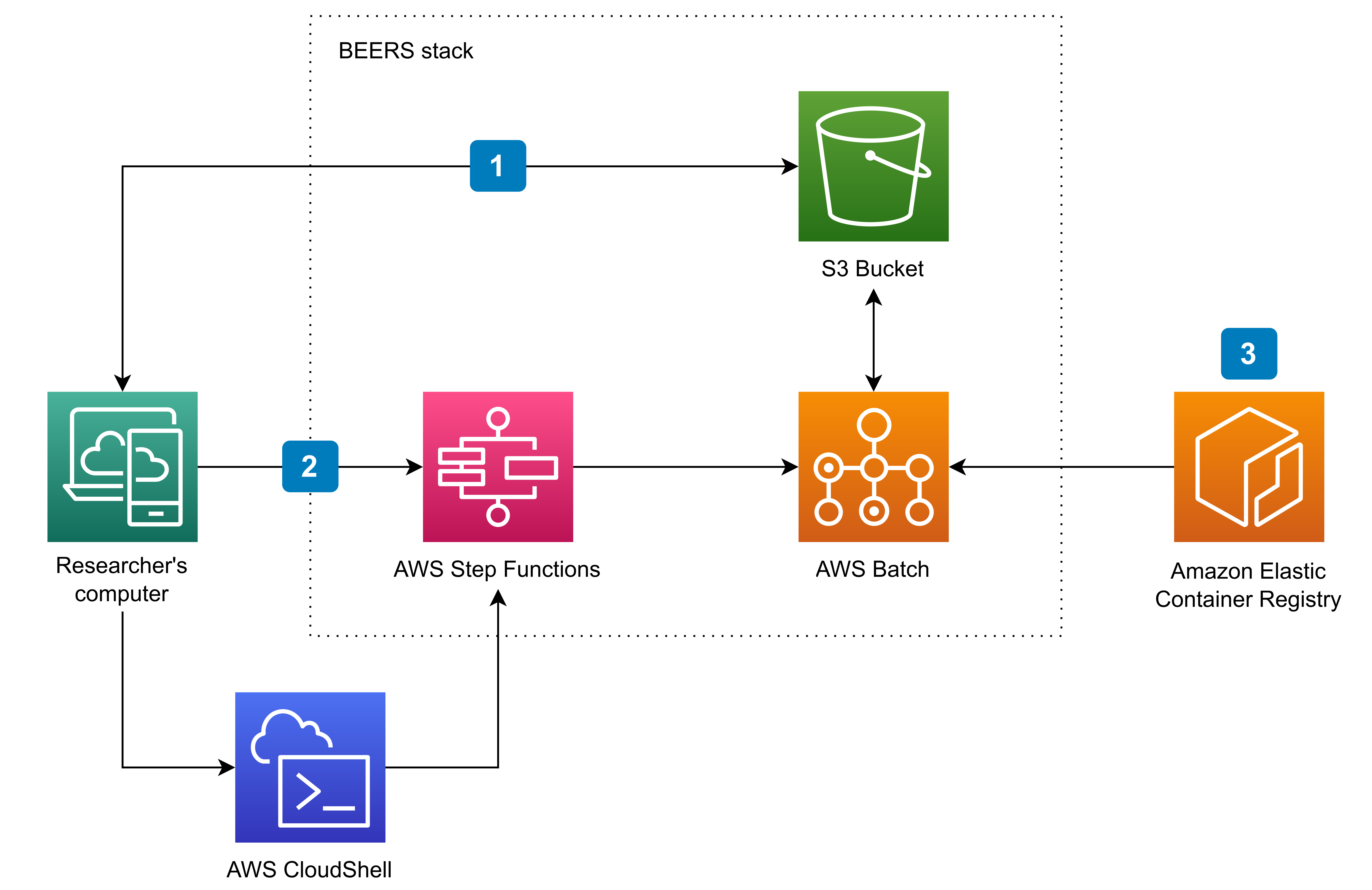

#### Figure S 1 - Running BEERS 2 in the Cloud

The BEERS stack encapsulates both the BEERS2 software, provided as an image in the container registry (3), as well as the hardware (cluster infrastructure powered by AWS Batch) necessary to run the software in a highly parallel fashion. Once the BEERS stack is deployed, researchers need to upload the necessary input data to the S3 bucket (1), and then start the pipeline execution by a press of a button in GUI (2) or an appropriate shell CLI command. The simulated data will be placed in the S3 bucket.

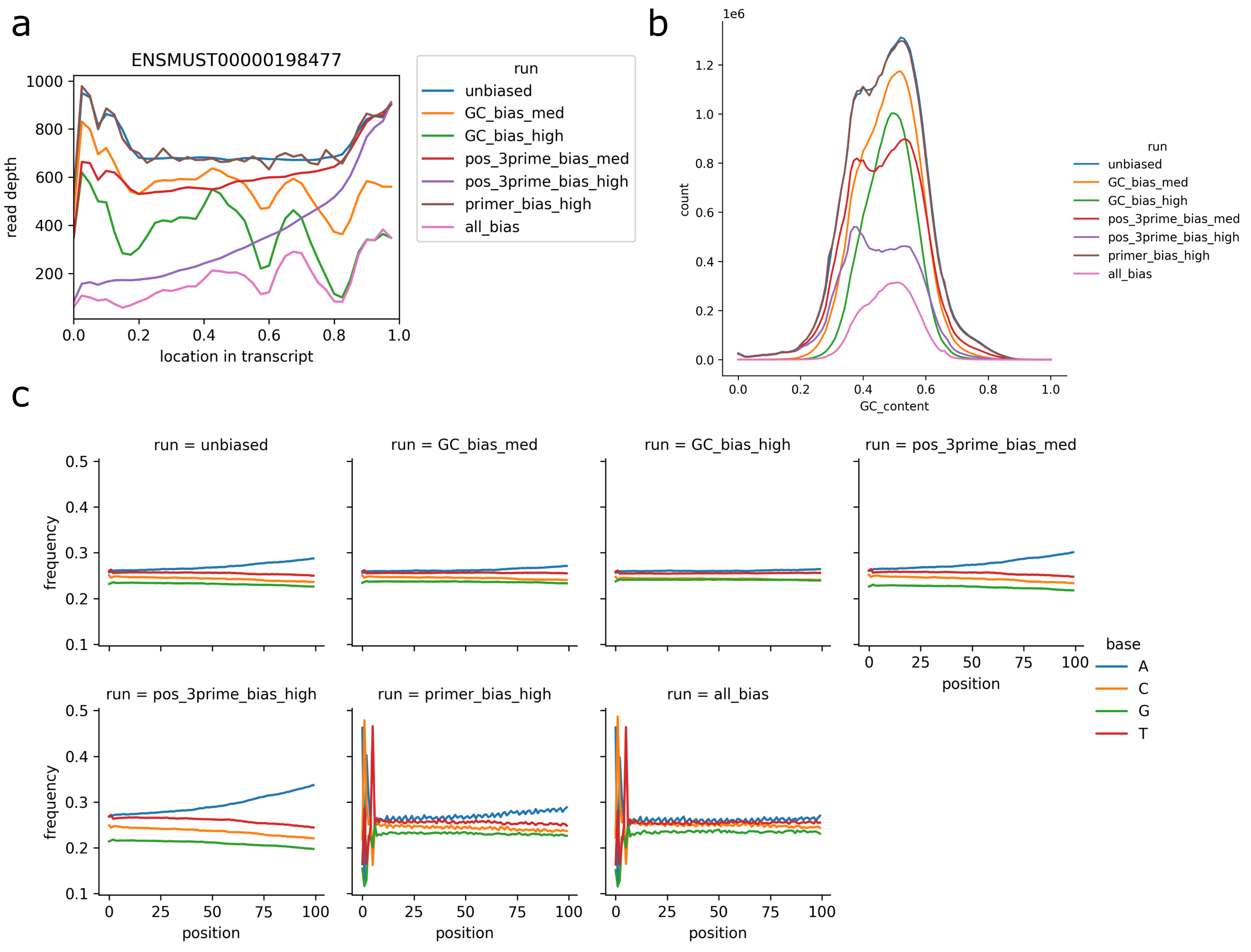

#### Figure S 2 - Bias effects in simulated data

We simulated the eight samples under varying bias configurations. For one representative sample, we plot the effects of those configurations on (a) coverage depth in one example transcript, (b) read GC content, and (c) fragment sequence bias of forward reads. Note that increasing quantities of A’s appear in the 3’ end of reads due to the presence of polyA tails in the data, though inclusion of GC or primer sequence bias reduces the amount of sequenced polyA tail.

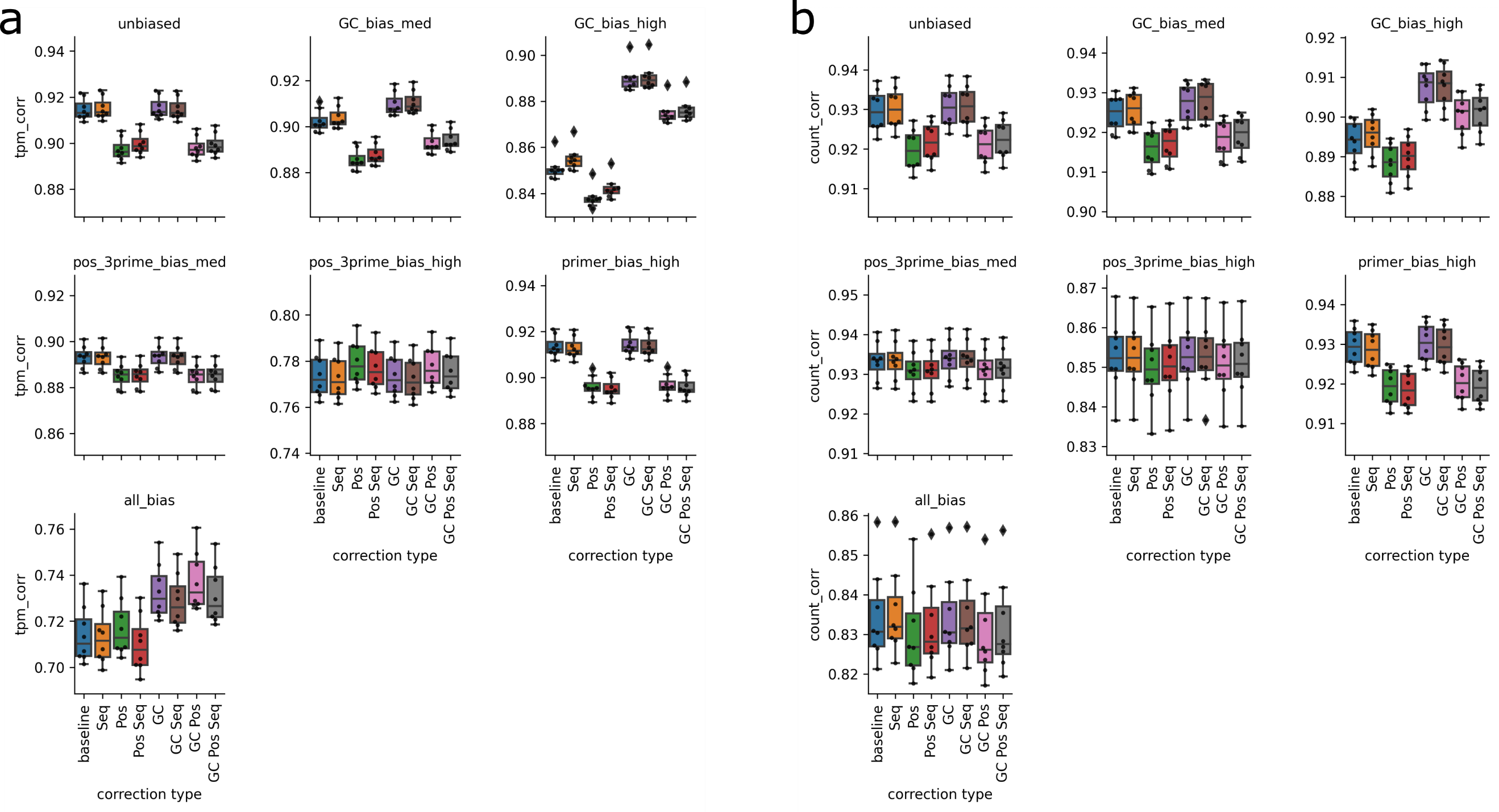

#### Figure S 3 – Correlation of true and quantified counts

For the eight samples simulated under seven different bias configurations, Salmon was run with each possible combination of bias correction options for fragment start/end sequence (Seq), position within transcript (Pos), and fragment GC content (GC). Spearman correlation of the true and estimated quantifications are plotted for (a) TPM values and (b) read count values. Boxes show quartiles.
